# Thermal adaptation constrains the temperature dependence of ecosystem metabolism

**DOI:** 10.1101/108696

**Authors:** Daniel Padfield, Chris Lowe, Angus Buckling, Richard Ffrench-Constant, Elisa Schaum, Simon Jennings, Felicity Shelley, Jón S. Ólafsson, Gabriel Yvon-Durocher

## Abstract

Gross primary production (GPP) is the largest flux in the carbon cycle, yet its response to global warming is highly uncertain. The temperature sensitivity of GPP is directly linked to photosynthetic physiology, but the response of GPP to warming over longer timescales could also be shaped by ecological and evolutionary processes that drive variation community structure and functional trait distributions. Here, we show that selection on photosynthetic traits within and across taxa dampen the effects of temperature on GPP across a catchment of geothermally heated streams. Autotrophs from cold streams had higher photosynthetic rates and after accounting for differences in biomass among sites, rates of ecosystem-level GPP were independent of temperature, despite a 20 ºC thermal gradient. Our results suggest that thermal adaptation constrains the long-term temperature dependence of GPP, and highlights the importance of considering physiological, ecological and evolutionary mechanisms when predicting how ecosystem-level processes respond to warming.

## INTRODUCTION

The carbon cycle is fundamentally metabolic (Falkowski *et al*. 2000). At the ecosystem level, gross primary production (GPP) represents the total amount of CO_2_ fixed by photosynthesis into organic carbon and is the largest flux in the global carbon cycle (Beer *et al*. 2010) transferring CO_2_ from the atmosphere to the biosphere, fuelling food webs and biological production (Field 1998). Understanding the mechanisms that shape how temperature influences rates of GPP across spatial, temporal and organisational scales is therefore an essential prerequisite to forecasting feedbacks between global warming and the carbon cycle.

Temperature can dictate rates of GPP over short timescales through its effects on photosynthetic physiology (Medlyn *et al*. 2002; Allen *et al*. 2005; Galmes *et al*. 2015). However, it is clear that over longer timescales (e.g. decades of gradual warming) ecological and evolutionary processes that mediate temperature induced changes in biomass, community composition and local adaptation of metabolic traits could feedback to influence the emergent effects of warming on ecosystem properties (Allen *et al*. 2005; Enquist *et al*. 2007; Michaletz *et al*. 2014; Cross *et al*. 2015). Indeed a recent analysis demonstrated that most of the variation in terrestrial primary production along a latitudinal temperature gradient could be explained by changes in biomass, and after controlling for biomass, rates were independent of temperature (Michaletz *et al*. 2014). Such temperature invariance in biomass-specific rates of primary production is counterintuitive considering the well-known exponential effects of temperature on the biochemistry of metabolism (Michaletz *et al*. 2014). Furthermore, it implies that selection on photosynthetic traits that compensate for the effects of temperature on physiological rates could play a fundamental role in mediating the effects of temperature on rates of primary production in the long-term (Kerkhoff *et al*. 2005; Enquist *et al*. 2007).

Here we investigate the interplay between the direct effects of temperature on photosynthesis, local adaptation through selection on photosynthetic traits, and changes in community biomass, on rates of gross primary production. We do so by extending the general model for ecosystem metabolism from metabolic theory (Enquist *et al*. 2003, 2007; Allen *et al*. 2005; Kerkhoff *et al*. 2005; Michaletz *et al*. 2014) to include the effects of thermal adaptation on the key traits that influence individual metabolism as well as potential temperature effects on ecosystem biomass pools. We then test our model’s predictions against empirical data collected from a catchment of naturally warmed Icelandic geothermal streams spanning a gradient of 20 ºC.

## THEORY

The metabolic theory of ecology (MTE) provides a powerful framework for understanding how temperature affects GPP by linking the photosynthetic rates of an ecosystem’s constituent individuals with the size and biomass structure of the community (Enquist *et al*. 2003, 2007; Allen *et al*. 2005; Kerkhoff *et al*. 2005; Yvon-Durocher & Allen 2012; Michaletz *et al*. 2014). Organism-level metabolism, *b*(*T*), responds predictably to temperature, increasing exponentially up to an optimum, followed by a more pronounced exponential decline (Fig. 1a). These thermal response curves can be quantified using a modification of the Sharpe-Schoolfield equation for high temperature inactivation (Schoolfield *et al*. 1981):

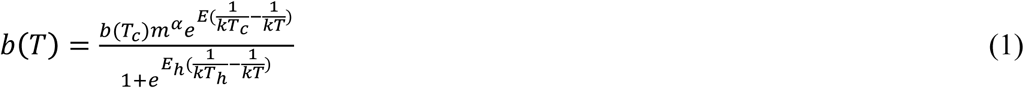

**Figure 1.**
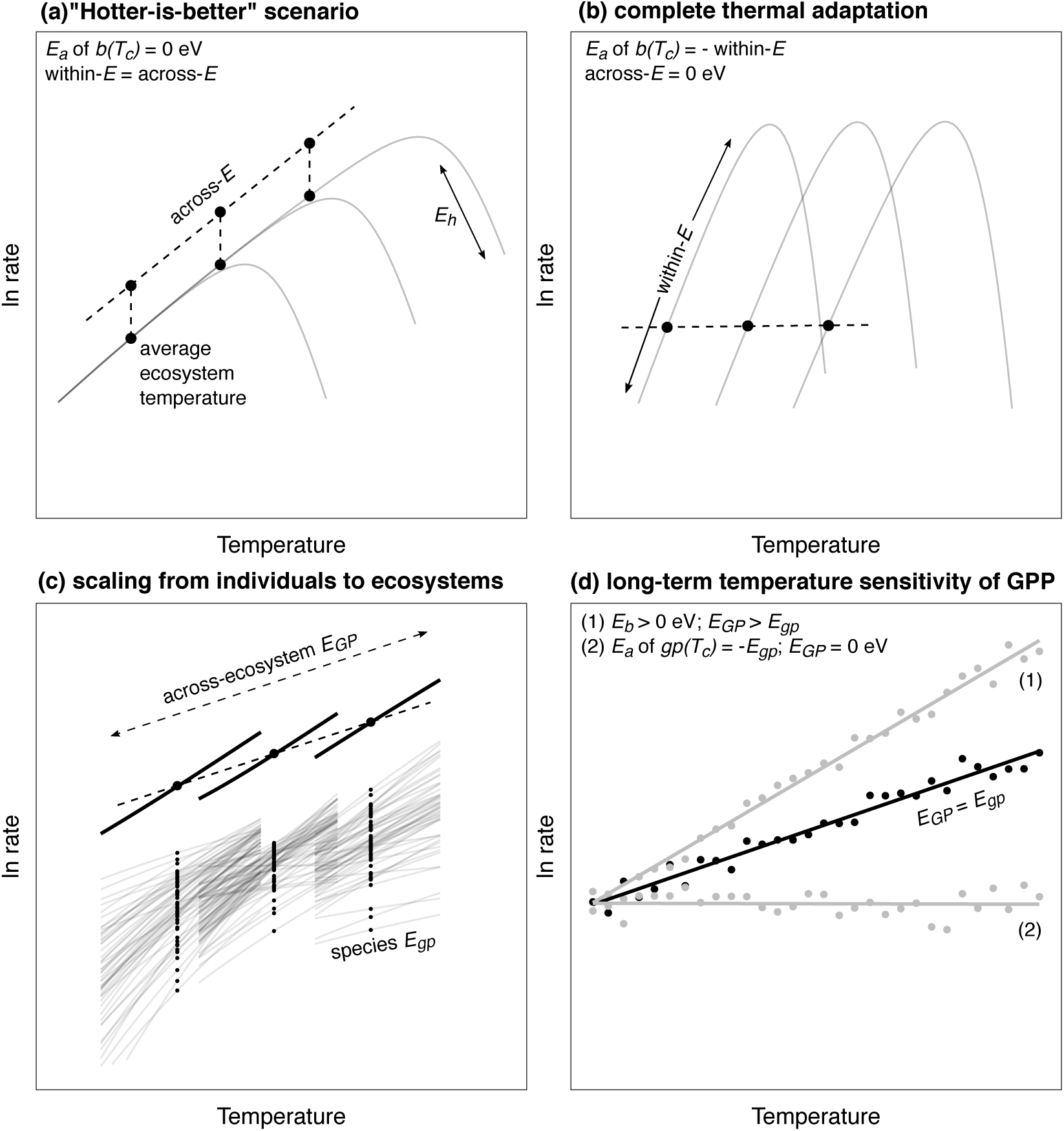
Scaling metabolism from organisms to ecosystems. (a) In a “hotter-is-better” scenario, thermodynamic constraints entirely dictate individual metabolic rates such that adaptation can only occur by moving peak performance up and down an “across-species” thermal performance curve. (b) Under complete thermal adaptation, an equalisation of peak rates occurs through upregulation of metabolic rates at cold, and downregulation of rates at high temperatures. (c) The long-term ecosystem temperature response, *E_GP_*, is an emergent property dependent on the thermal response of each ecosystem’s constituent individuals. (d) If local thermal adaptation drives temperature dependence in the metabolic normalisation (e.g. as expected under the ‘complete thermal adaptation’ hypothesis) or standing biomass is temperature dependent, the long-term temperature sensitivity of ecosystem metabolism may deviate away from the average activation energy of individual metabolism.

Where *b*(*T*) is the rate of metabolism at temperature *T*, in Kelvin (K), *k* is Boltzmann’s constant (8.62 × 10^−5^ eV K^-1^), *E* is the activation energy (in eV), *E_h_* characterises temperature-induced inactivation of enzyme kinetics above*T_h_*, which is the temperature at which half the enzymes are inactivated. In this expression, *b*(*T_c_*) is the rate of metabolism normalised to a reference temperature (e.g. 10 ºC), where no low or high temperature inactivation occurs and *m^α^* is the mass dependence of metabolic rate characterised by an exponent *α,* that ranges between ¾ and 1 across multicellular and unicellular autotrophs (Gillooly *et al*. 2001; DeLong *et al*. 2010). Equation 1 yields a maximum metabolic rate at an optimum temperature,

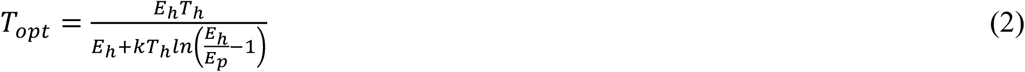

The parameters in equations 1 & 2, which govern the height and shape of the thermal response curve can be considered “metabolic traits” (Padfield *et al*. 2016) and have long been known to shift as organisms adapt to new thermal environments (Berry & Bjorkman 1980; Huey & Kingsolver 1989). Equation 1 can be simplified to the Arrhenius equation,

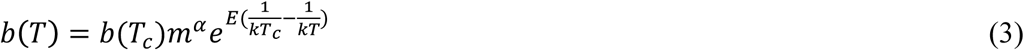

which captures only the rising part of the thermal response curve, if the temperatures organisms experience in the environment are below *T_opt_* (Savage *et al*. 2004; Dell *et al*. 2011; Sunday *et al*. 2012). We use this simpler, more tractable model of the temperature dependence in the following theory, which attempts to explore the mechanisms driving the emergent temperature sensitivity of ecosystem-level gross primary production.

At the organism-level, the size and temperature dependence of gross photosynthesis can be characterized as:

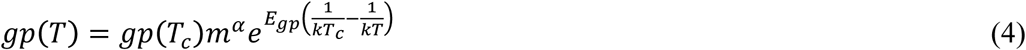

where *gp*(*T*) is the rate of gross photosynthesis and temperature *T*, and *gp*(*T_c_*) is the rate of gross photosynthesis normalised to a reference temperature and *E_gp_* is the activation energy of gross photosynthesis. Net photosynthesis, *np*, which is the amount of photosynthate available for allocation to biomass production after accounting for autotroph respiration is given by,

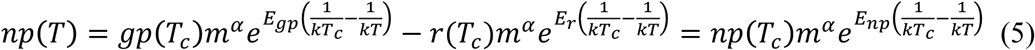

where *np*(*T*) is the rate of net photosynthesis at temperature *T*, *r*(*T_c_*) is the rate of respiration normalised to a reference temperature, *T_c_*, and *E_np_* and *E_r_* are the activation energies of net photosynthesis and respiration. The form of equation 5 implies that the temperature sensitivity of *np* will not strictly follow a simple Boltzmann-Arrhenius relation (see supplementary information for a derivation of *E_np_*). Nevertheless, we can approximate the temperature sensitivity of net photosynthesis using an apparent activation energy, *E_np_*, with a reasonable degree of accuracy (Fig. S7).

Using Equation 4 and principles from MTE, the rate of gross primary productivity per unit area, *A*, can be approximated by the sum of the photosynthetic rates of its constituent organisms (Fig. 1c):

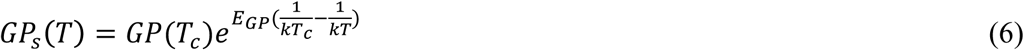

where *GP_s_*(*T*) is the rate of gross primary production in ecosystem *s*, at temperature *T*, 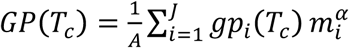, is the ecosystem-level metabolic normalisation, where *J* is the total number of individual organisms, *i*, which comprise all autotrophs in *s*. In equation 6, the apparent long-term temperature dependence of gross primary production, *E_GP_*, is assumed to be equal to that of the average temperature dependence for individual-level gross photosynthesis, *E_gp_*, provided that the ecosystem-level normalisation, *GP*(*T_c_*) is independent of temperature (Fig. 1d). However, if *gp_i_*(*T_c_*) or total biomass, 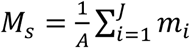, exhibit temperature dependence, for example via temperature driven selection on *gp_i_*(*T_c_*) or covariance between resource availability, temperature and *M_s_*, then the scaling of the activation energy from individuals to ecosystems will no longer hold (e.g. *E_GP_* ≠ *E_gp_*). Thus, ecological processes that influence *M_S_* and evolutionary dynamics which shape variation in *gp_i_*(*T_c_*) have the potential to play an integral, but as yet underappreciated role in mediating the response of ecosystem metabolism to temperature if they modify the metabolic capacity of ecosystem biomass pools (but see Kerkhoff *et al*. 2005; Enquist *et al*. 2007; Michaletz *et al*. 2014).

Previous work on aquatic and terrestrial autotrophs has shown that autotrophs adapt to long-term temperature changes by shifts in the respiratory and photosynthetic normalisation constant; up-regulating rates at low temperatures and down-regulating at high temperature, to alleviate the constraints of thermodynamics on enzyme kinetics (Atkin *et al*. 2015; Padfield *et al*. 2016; Reich *et al*. 2016; Scafaro *et al*. 2016). We therefore expect *gp_i_*(*T_c_*) to exhibit temperature dependence along long-term thermal gradients, which in the absence of an explicit first principles derivation, we can approximate as

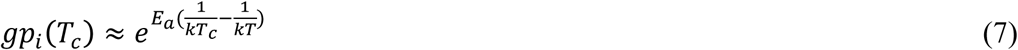

where *E_a_* is an adaptation parameter that characterises the change in *gp_i_*(*T_c_*) with temperature owing to thermal adaptation. Substituting the temperature dependence for *gp_i_*(*T_c_*) into equation 6 and simplifying, yields the following expression for the temperature dependence of gross primary production,

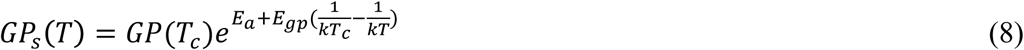

Under the “hotter-is-better” model of thermal adaptation (Fig. 1a), where a single activation energy governs the temperature dependence of metabolism within and across species (Gillooly *et al*. 2001; Savage *et al*. 2004; Angilletta *et al*. 2010) and *E_a_* = 0, the ecosystem-level activation energy would equal that of individual-level metabolism (i.e. *E_GP_* = *E_gp_*; Fig. 1d) – this is the typical assumption made in metabolic theory (Brown *et al*. 2004; Demars *et al*. 2016). However, when *E_a_* ≠ 0, *E_GP_* = *E_a_* + *E_gp_*, and the ecosystem-level activation energy will deviate from the average organism-level temperature dependence owing to the effects of thermal adaptation on *gp_i_*(*T_c_*). If thermal adaptation results in complete compensation (i.e. *E_a_* = − *E_gp_*; Fig. 1b), and *M_s_* does not covary with temperature, then ecosystem” level gross primary production will be independent of temperature (i.e. *E_GP_* = 0; Fig. 1d). Following the same reasoning, any temperature dependence in *M_s_* will also result in deviations from the average individual-level activation energy. For example, recent experimental work has shown that covariance between temperature and rates of nutrient cycling can cause *M_s_* to increase with temperature (Welter *et al*. 2015; Williamson *et al*. 2016)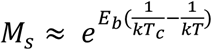, where *E_b_* is the activation energy characterising the temperature dependence of total biomass. When *E_b_* > 0, substituting in the temperature dependence for *M_s_* into equation 8 leads to an increase in the ecosystem-level activation energy regardless of the mode of thermal adaptation (*E_GP_* = *E_gp_* + *E_b_* + *E_a_*; Fig. 1d). This model emphasises how different ecological and evolutionary mechanisms that drive temperature dependent variation in individual-level metabolic traits and/or ecosystem biomass pools can influence the emergent long-term temperature sensitivity ecosystem metabolism (Fig. 1c:d).

We now use measurements of the temperature dependence of organism” and ecosystem-level photosynthesis from a catchment of naturally warmed geothermal streams to test the expectations of our model and investigate how ecological and evolutionary processes shape the long-term temperature sensitivity of GPP. Critically, this system allows us to measure photosynthetic responses to temperature at both organism and ecosystem scales from sites that are in close proximity, yet differ substantially in their thermal history (i.e. 20 ºC *in situ* temperature gradient among sites).

## METHODS

### Study site

The study was conducted in a geothermally active valley close to Hveragerdi village, 45 km east of Reykjavik, Iceland. The area contains a large number of mainly groundwater-fed streams that are subjected to differential natural geothermal warming from the bedrock (O’Gorman *et al*. 2014). Twelve streams have been mapped in the valley with average temperatures ranging from 7 – 27 ºC (Fig. S1 & Table S1). We measured a number of physical (width, depth, velocity) and chemical (pH, conductivity, nitrate, nitrite, soluble reactive phosphate, ammonium) variables across the stream catchment (Table S2) and none of these variables were significantly correlated with temperature (Table S3). The study was carried out during May and June in 2015 and 2016.

### Measuring the population level metabolic thermal response

We sampled 13 of the most abundant autotrophic biofilm taxa from 8 streams spanning the catchment’s full thermal gradient. Multiple taxa were removed from four streams where more than one taxon was at high density (Table S4). Measurements first entailed characterising a photosynthesis-irradiance (PI) curve from 0 – 2000 µmol m^-2^ s^-1^ at the average stream temperature for each taxon. Net photosynthesis (*np*) was measured as O_2_ evolution in a Clark-type oxygen electrode (Hansatech Ltd, King's Lynn UK Chlorolab2) at increasing light intensities in intervals of 50 µmol^-1^ m^-2^ s^-1^ up to 300 µmol^-1^ m^-2^ s^-1^, and then in intervals of 100 µmol^-1^ m^-2^ s^-1^ up to 1000 µmol^-1^ m^-2^ s^-1^, followed by 200 µmol steps up to 2000 µmol^-1^ m^-2^ s ^-1^. Rates of respiration (*r*) were measured as O_2_ consumption in the dark. This yielded a PI curve from which the optimal light intensity for net photosynthesis was estimated using a modification of Eilers’ photoinhibition model (Eilers & Peeters 1988) fitted via nonlinear least squares regression (Fig. S2):

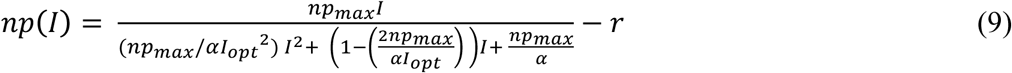

where *np*(*I*), is the rate of net photosynthesis at irradiance, *I*, *np*_*max*_ is the photosynthetic maximum that occurs at optimal light, *I*_*opt*_, *α* controls the gradient of the initial slope and *r* is respiration, the rate of oxygen consumption in the dark. The optimum light intensity (*I*_*opt*_, μmol^−1^ m^−2^ s^−1^) for each taxon was then used for measuring net photosynthesis at all other assay temperatures in the acute thermal gradient experiments. Rates of gross photosynthesis were calculated by the summation of the measured rates of net photosynthesis and respiration.

Rates of photosynthesis and respiration were normalised to biomass by expressing rates per unit of chlorophyll *a*. Chlorophyll *a* extraction was achieved by grinding the sample tissue with methanol for 5 minutes, centrifugation and measuring chlorophyll *a* extinction coefficients on a spectrophotometer. Total chlorophyll *a* (μg) was then calculated by measuring absorbance at 750 nm, 665 nm and 632 nm.

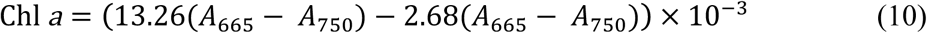

Acute temperature responses of biomass normalised gross and net photosynthesis and respiration were fitted to the modified Sharpe-Schoolfield equation for high temperature inactivation (Equation 1). Best fits for each thermal response curve were determined using non-linear least squares regression using the ‘nlsLM’ function in the ‘minpack.lm’ (Elzhov *et al*. 2009) package in R statistical software (R Core Team 2014; v3.2.2), following the methods outlined in Padfield *et al*., (2016).

We tested for thermal adaptation by assessing whether the parameters in eqns. 1 and 2 as well as the rate of gross photosynthesis at the average stream temperature, *gp*(*T_s_*) varied systematically with stream temperature. We fitted the metabolic traits to a modified Boltzmann-Arrhenius function within a linear mixed effects modelling framework:

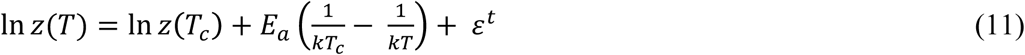
where *z* is the metabolic trait at stream temperature, *T*, *z*(*T_c_*) is the value of the trait at the mean temperature across all streams, *T_c_*, and *E_a_* is the activation energy that determines how much *z* changes as a function of *T* due to thermal adaptation and *ɛ^t^* is a random effect on the intercept accounting for multiple measurements of the same metabolic trait of each isolated biofilm taxon (i.e. one value each for gross and net photosynthesis and respiration). We fitted eq. 11 to each metabolic trait with stream temperature, flux (3 level factor with ‘gross’ and ‘net photosynthesis’ and ‘respiration’) and their interaction as fixed effects (Table S5). Significance of the parameters were determined using likelihood ratio tests. Model selection was carried out on models fitted using maximum likelihood and the most parsimonious model was refitted using restricted maximum likelihood for parameter estimation.

### Measuring *in situ* rates of ecosystem-level gross primary production

Ecosystem metabolism was calculated from measurements of dissolved oxygen over time using the single station method (Odum 1956). Sensors were deployed in all streams and at multiple sites within a stream where temperature gradients existed within streams due to differential geothermal warming. Dissolved oxygen concentration and temperature were monitored at 1-minute intervals using miniDOT dissolved oxygen loggers (PME Inc) (Fig. S3 & Fig. S5). Light sensors (LI-COR Inc) were deployed simultaneously at two sites in the centre of the catchment. Physical variables of each stream, including the depth (m), width (m), velocity (m s^-1^), were measured along horizontal transects at 10 m intervals up to 100 m upstream of the sensor deployment. Values for depth, width and velocity were averaged across the reach (Table S2).

The change in O_2_ concentration at a single station between two subsequent measurements (∆*DO*) can be approximated as:

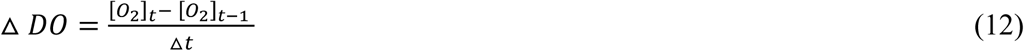

with [O_2_]_t_ the concentration of oxygen (mg L^-1^) at time *t* and can be modelled using a framework based on the Odum’s O_2_ change technique (Odum 1956):

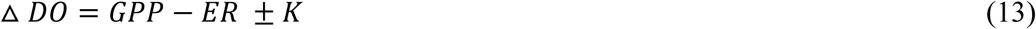

where *GPP* (g m^-3^ hr ^-1^) is the composite of volumetric gross primary productivity, minus volumetric ecosystem respiration, *ER* (g m^-3^ hr ^-1^) and *K* is the net exchange of oxygen with the atmosphere (g O_2_ m^-3^). The net exchange of oxygen with the atmosphere is the product of the O_2_ gas transfer velocity, *k* (m min^-1^), and the O_2_ concentration gradient between the water body and the atmosphere (temperature and atmosphere corrected DO concentration at 100% saturation minus [O_2_]_t_) over the measurement interval.

The gas transfer velocity, *k* (m min^-1^), was calculated using the surface-renewal model and corrected for the stream temperature:

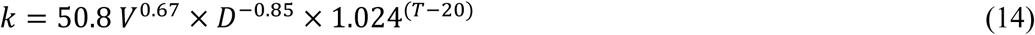

where *V* is velocity (cm s^-1^), *D* is the mean stream depth (cm) adjusted for stream temperature, *T* (Bott 1996). This value was subsequently transformed into (m h^-1^). Estimated rates of reaeration, derived using the surface renewal model from measurements of velocity and depth, correspond well to reaeration rates measured experimentally using propane additions in an adjacent Icelandic catchment with comparable physico-chemical characteristics (see Fig. S8; Demars *et al*. 2011).

The net metabolic flux for a given measurement interval is equal to △ *DO* – *K*. During the night (where light < 5 µmol m^-2^ s^-1^), GPP is zero, so the net metabolic flux is equal to ER. During the day, ER was determined by interpolating average ER over the defined night period. GPP for each daytime interval was the difference between net metabolism flux and interpolated ER. Daily volumetric rates of GPP (g O_2_ m^-3^ day^-1^) were calculated as the sum of the 15-minute rates over each 24-hour period. Volumetric rates were converted to areal units (g O_2_ m^-2^ day^-1^) by multiplying by the mean water depth of the stream reach.

We measured autotrophic biomass density (g Chl *a* m^-2^) across the stream catchment by taking measurements of chlorophyll *a*. A core of 28.27 cm^2^ was removed from 3 randomly chosen rocks and chlorophyll *a* was measured using the extraction protocol detailed above. The total standing biomass, *M_s_*, of each stream reach was estimated by multiplying average biomass density by the total reach area, which was estimated from the mean width and the distance upstream from the oxygen sensor integrated over (Chapra & Di Toro 1991; Demars *et al*. 2015),

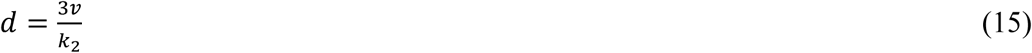

where three times the velocity of the stream (*v*; m d^-1^) divided by the gas transfer coefficient (*K*
_
*2*
_; d^-1^) gives the approximation of the distance upstream integrated by the single station method (*d*; m) (Grace & Imberger 2006). Biomass normalised rates of GPP per stream (g O_2_ g Chl *a*
^-1^ day^-1^) were calculated by dividing areal rates of GPP by the total standing biomass in the upstream reach.

We used linear mixed-effects modelling to investigate the temperature dependence of GPP across catchment, allowing us to control for the hierarchical structure of the data (e.g. variance of days nested within years nested within streams). We characterised the temperature dependence of GPP with a linearised version of the Boltzmann-Arrhenius function in a linear mixed effects model:

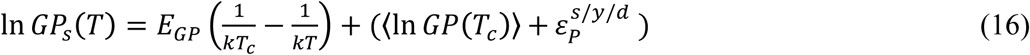
where *Gp_s_*(*T*) is the rate of gross primary production in stream *s* on year *y* on day *d* at temperature *T* (Kelvin), *E_GP_* is the activation energy (eV) which characterises the exponential temperature sensitivity of photosynthetic rates, 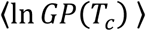 is the average rate of *GP* across streams and days normalised to *T_c_* = 283 K (10 °C) and 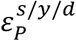 is a nested random effect that characterises deviations from 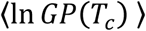 at the level of *d* within *y* within *s*. Significance of the parameters and model selection was carried out as described above for the analysis of the population-level metabolic traits (Table 1).

**Table 1.** Results of the linear mixed effects model analysis for gross primary productivity (GPP) for all years and 2016 only. The results of the model selection procedure on the fixed effect terms are given and the most parsimonious models are highlighted in bold. Analyses reveal that GPP increased significantly with stream temperature. The analyses for 2016 show that the observed temperature response was driven by covariance between biomass and temperature rather than the direct effects of temperature on rates of photosynthesis *per se*.

| Model | d.f. | AICc | log Lik | L-ratio | P |
| --- | --- | --- | --- | --- | --- |
| **All years :** |  |  |  |  |  |
| random effects structure |  |  |  |  |  |
| random = 1 \| stream/year/day |  |  |  |  |  |
| fixed effects structure |  |  |  |  |  |
| 1. **ln GPP ~ 1 + stream temperature** | **6** | **82.9** | **-34.0** |  |  |
| 2. ln flux ~ 1 | 5 | 85.8 | -36.9 | 5.80 | **0.016** |
| **2016 only :** |  |  |  |  |  |
| random effects structure |  |  |  |  |  |
| random = 1 \| stream/day |  |  |  |  |  |
| fixed effects structure |  |  |  |  |  |
| 1. ln GPP ~ 1 + stream temperature + biomass | 6 | 48.8 | -14.9 |  |  |
| 2. **ln GPP ~ 1 + biomass** | **5** | **45.3** | **-15.3** | 0.87 | 0.35 |
| 3. ln flux ~ 1 | 4 | 45.8 | -17.4 | 4.25 | **0.04** |

We tested for the effect of total biomass and temperature on GPP across the catchment using the data from 2016 (where we also quantified autotroph biomass) by undertaking a multiple regression by expanding eq. 16 to include the effect the biomass on GPP:

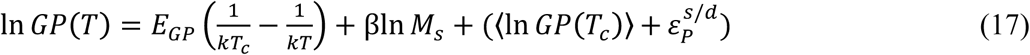

where β characterises the power-law scaling of *GP*(*T*) with *M_s_* and the random effects specification changed to account for deviation from 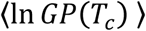 between days nested within streams. Model selection was as described above (Table 1).

### Inorganic nutrients

Water samples for measuring dissolved inorganic nutrient concentrations (NO_2_, NO_3_, NH_4_ and PO_4_; µmol L^-1^) were collected from each stream. Samples were filtered (Whatmann GF/F) and stored frozen at -20 ºC for subsequent analysis using a segmented flow auto-analyser (Table S3) (Kirkwood 1996).

## RESULTS

### Population level metabolism

Macroscopic cyanobacteria, filamentous eukaryotic algae, and bryophytes were the dominant autotrophs across the catchment (Table S4). To investigate how long-term differences in temperature shaped variation in photosynthetic traits across the catchment, we sampled the most abundant autotroph taxa from 8 streams spanning the full temperature gradient and measured the acute responses of gross photosynthesis and respiration to temperatures spanning 5 to 50 ºC. Gross photosynthesis and respiration followed unimodal responses to acute temperature variation and were well fit by equation 1 (Fig. 2a-b). We predicted exponential declines in the metabolic normalisation constants, moving from cold to warm environments, owing to the effects of thermal adaptation. Consistent with this hypothesis, the log-transformed rates of gross photosynthesis, (ln *gp*(*T_c_*)) and respiration (ln *r*(*T_c_*)) normalised to a reference temperature, *T_c_* = 10 ºC, declined linearly with increasing stream temperature with the same activation energy (*E_a_* = -0.64 eV; 95% CI: -1.22 to -0.05 eV; Fig. 2c).Since *np*(*T_c_*) = *gp*(*T_c_*) − *r*(*T_c_*), the normalization for net photosynthesis also declined with increasing temperature with an *E_a_* = -0.64 eV.

**Figure 2.**
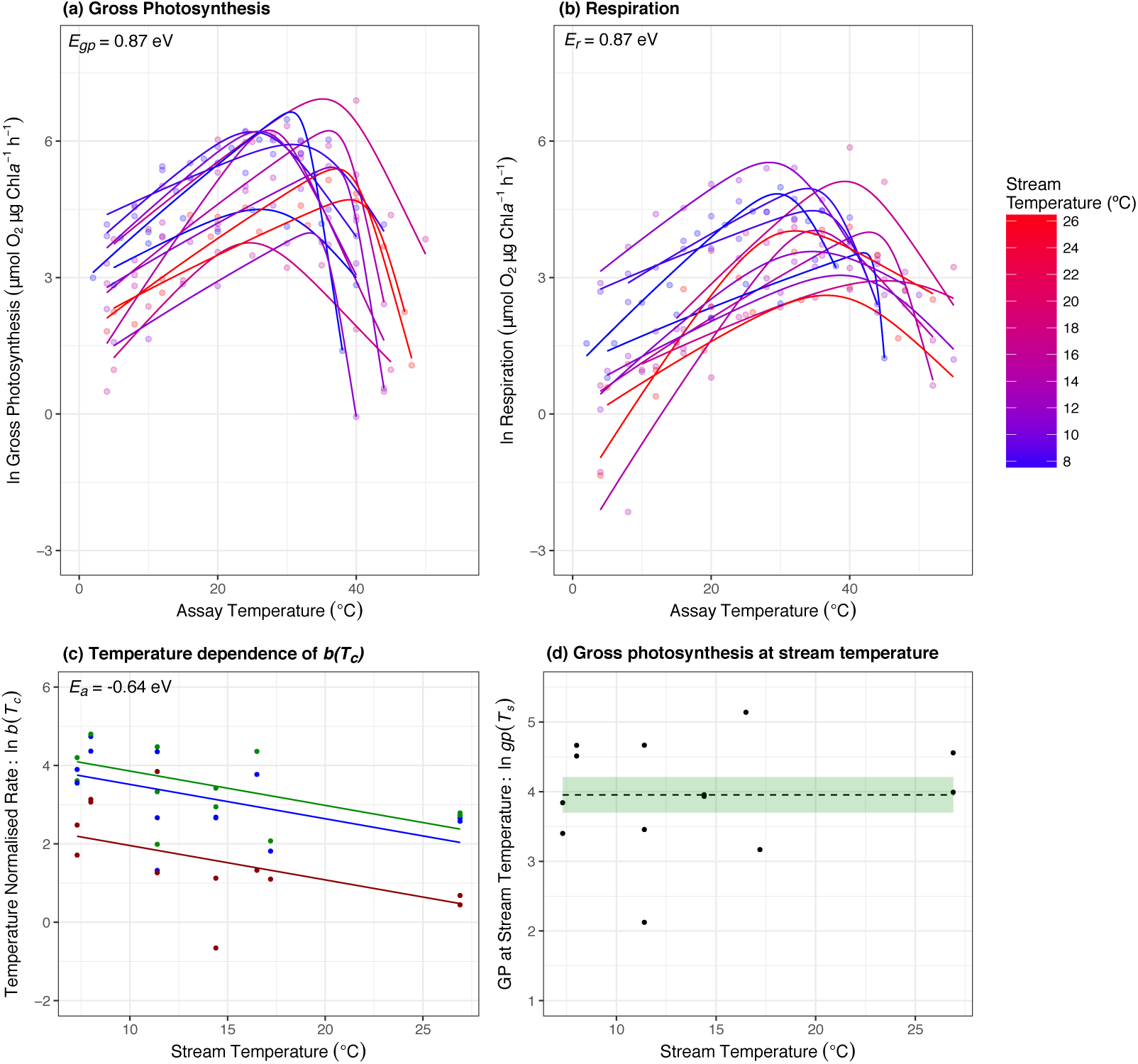
Patterns of metabolic thermal adaptation. (a, b) Acute thermal response curves for gross photosynthesis and respiration were measured for each isolated autotroph from streams spanning average temperatures from 7 ºC (blue) to 27 ºC (red). Fitted lines are based on the best-fit parameters from non-linear least squares regression using the modified Sharpe-Schoolfield model (see Methods). (c) Metabolic rates normalised to 10 ºC, *b*(*T_c_*), decrease exponentially with increasing stream temperature for gross photosynthesis (green), net photosynthesis (blue) and respiration (red) (d) Rates of gross photosynthesis at the average stream temperature showed no temperature dependence. Fitted lines and coloured bands in (c) and (d) represent the best fit and the uncertainty of the fixed effects of the best linear mixed effect model.

Because the dominant autotroph taxa varied across the streams (Table S4), the decline in the photosynthetic trait, *gp*(*T_c_*), with increasing stream temperature is likely influenced by species sorting (e.g. filtering of species and traits from the regional species pool). To investigate whether adaptive evolution also played a role, we analysed data from only the most common genera *Nostoc,* which was distributed across 5 streams spanning a gradient of 10.2 ºC. *gp*(*T_c_*), *np*(*T_c_*) and *r*(*T_c_*) also decreased with increasing stream temperature in *Nostoc* with the thermal sensitivity not significantly different from that of all the autotroph taxa together (Fig. S6). This trend provides evidence for local thermal adaptation. An important consequence of the decrease in *gp*(*T_c_*) with increasing stream temperature was that rates of gross photosynthesis at the average temperature of each stream, *gp*(*T_s_*), were independent of temperature (Fig. 2d), indicating that species sorting and adaptation led to complete compensation of organism-level metabolism over the catchment’s thermal gradient.

Both the optimum temperature, *T_opt_*, and *T_h_*, which is the temperature at which half the enzymes are inactivated, were positively correlated with average stream temperature (Table S5) providing further evidence for local adaptation. We found no evidence for systematic variation in the activation or inactivation energies (*E_a_* or *E_h_*) across the thermal suggesting these traits are unlikely to be under temperature dependent-selection (Table S5). Previous work has often shown that photosynthesis has a lower activation energy than respiration (Allen *et al*. 2005; López-Urrutia *et al*. 2006; Padfield *et al*. 2016). In contrast, we found that the average activation energies of gross photosynthesis and respiration were not significantly different and could be characterised by a common activation energy, *E* = 0.87 eV; 95% CI = 0.77 to 0.97 eV. Similarly, *E_h_*, which characterises inactivation of kinetics past the optimum was not significantly different between fluxes and could be characterised by a common value for respiration and photosynthesis (*E_h_* = 4.91 eV; 95% CI: 3.95 – 5.97 eV).

### Ecosystem level gross primary productivity

Based on the observation that the activation energies of gross photosynthesis and the adaptation parameter were of equal magnitude and opposite sign, our model for the scaling of metabolism from organisms to ecosystems (Eq. 8) predicts that rates of gross primary production should be independent of temperature across the catchment (e.g. *E_GP_* = *E_gp_* + *E_a_* ≈ 0), provided that biomass does not covary with temperature. We measured rates of *in situ* GPP in 11 streams across the catchment’s full temperature gradient in 2015 and 2016. Rates of GPP increased with average stream temperature and the long-term temperature sensitivity of GPP (characterised by fitting the Boltzmann-Arrhenius function [see Methods]) yielded an activation energy of *E_GP_* = 0.57 eV (95% CI: 0.10 − 1.04 eV; Fig. 3a).

To investigate potential covariance between temperature and biomass, *M_s_*, and its impact on the temperature dependence of GPP, we also quantified *in situ* standing autotrophic biomass. Autotroph biomass density, *M_s_*, increased systematically with temperature across the catchment with a temperature sensitivity of *E_b_* = 0.68 eV (95% CI: 0.24 − 1.12 eV; Fig. 3b). The similarity between *E_GP_* and *E_b_* – they have 95% confidence intervals that overlap – indicates that covariance between biomass and temperature could be the main driver of the temperature dependence of GPP across the catchment.

We quantified the effects of both temperature and *M_s_* on GPP using multiple regression in a mixed effects modelling framework (see Methods). The best fitting model included only ln (*M_s_*) as a predictor (Table 1; Fig. 3c) and after controlling for variation in ln (M_s_), rates of GPP were independent of temperature across the catchment (Table 1; Fig. 3d). These findings are consistent with predictions from our model and provide evidence that systematic variation in the photosynthetic normalisation owing to thermal adaptation results in complete compensation of biomass-specific metabolic rates at organism and ecosystem scales.

**Figure 3.**
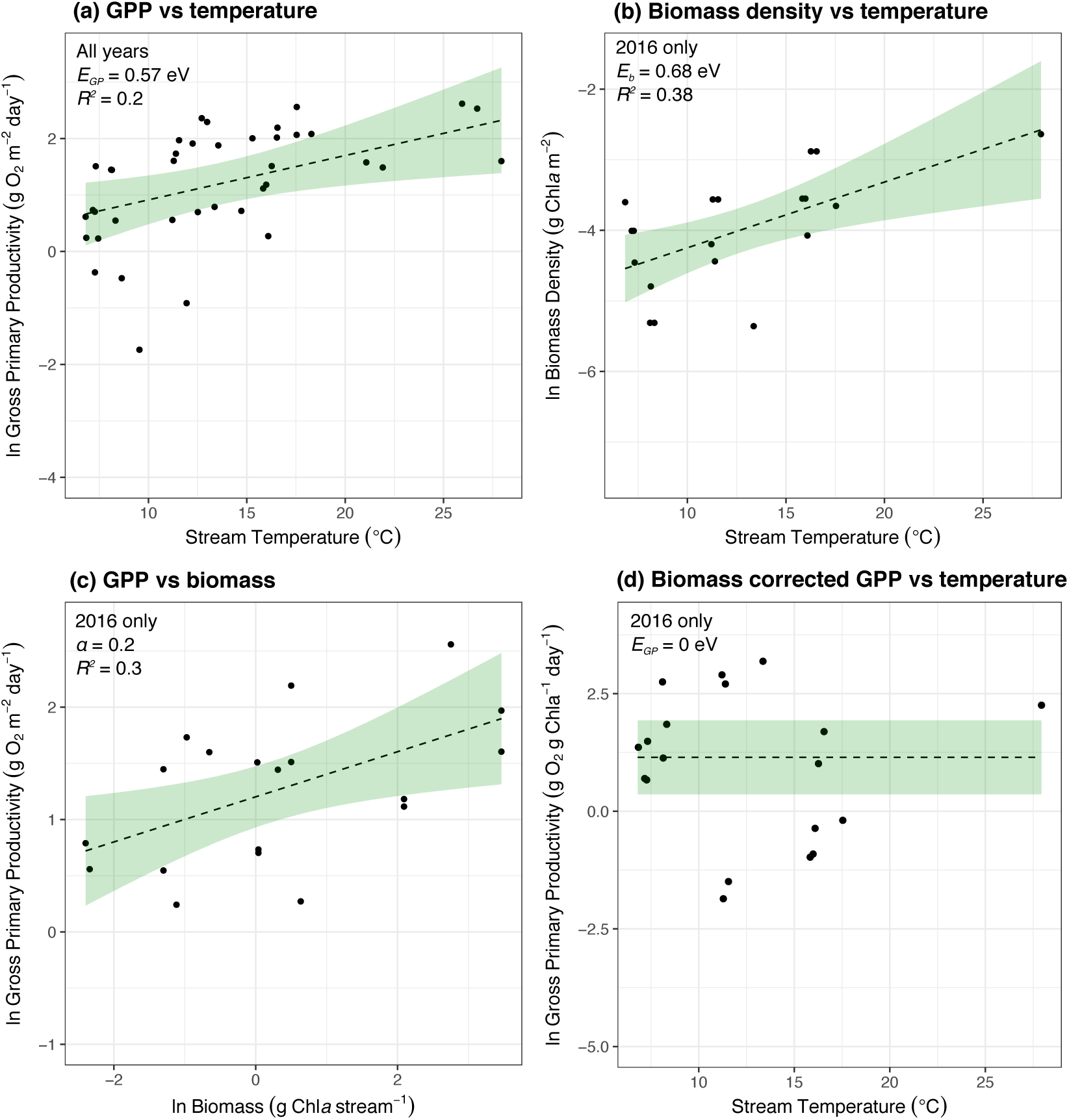
The effects of temperature and biomass on gross primary productivity. Gross primary productivity (a) and biomass density (b) increase with temperature across the catchment. (c) A multiple regression shows that variation in GPP is driven primarily by changes in biomass. (d) After accounting for biomass, rates of GPP are invariant with respect to temperature across the catchment. Fitted lines in (a, c, d) represent the best fit and the uncertainty of the fixed effects of the best linear mixed effect model (Table 1). In (b) the lines represent the fitted line and associated confidence interval of a linear regression.

## Discussion

Understanding how ecosystem-level properties like gross primary production (GPP) will respond to global warming is of central importance to predicting the response of the carbon cycle and contributing biogeochemical and food web processes to climate change. It is however a major challenge that requires an integration of physiological, ecological and evolutionary processes that together shape the emergent response of ecosystem metabolism to long-term changes in temperature. We have addressed this key problem by extending the general model of ecosystem metabolism from metabolic theory (Enquist *et al*. 2003, 2007; Allen *et al*. 2005; Kerkhoff *et al*. 2005) and testing its predictions at organism and ecosystem scales in a catchment of naturally warmed geothermal streams. Our model and analyses demonstrate that temperature-dependent selection on organism-level metabolic traits and shifts in ecosystem biomass can be as important as the direct effects of temperature on metabolism in shaping the temperature dependence of GPP.

Our model predicted that when the temperature dependence of the metabolic normalisation constant across taxa inhabiting environments with different thermal histories is of a similar magnitude but opposing sign to that of organism-level metabolism, the two temperature sensitivities cancel, rendering biomass-specific metabolic rates independent of temperature. Measurements of the thermal response curves for photosynthesis and respiration from the autotrophs isolated across the 20 ºC *in situ* gradient provided strong support for this prediction, with rates of gross photosynthesis invariant with respect to differences in average *in situ* temperatures and activation energies of organism-level gross photosynthesis and the photosynthetic normalisation, *gp*(*T_c_*), across taxa that were not significantly different and of opposite sign.

The exponential decline in *gp*(*T_c_*) along the *in situ* thermal gradient primarily reflected turnover in the composition of the dominant autotroph taxa across the streams driven by species sorting. This result is in line with work demonstrating declines in the metabolic normalisation constant across vascular plant species along broad-scale latitudinal gradients in terrestrial ecosystems (Atkin *et al*. 2015). However, we also found a comparable negative temperature dependence of *gp*(*T_c_*) in the genera, *Nostoc*, which was distributed across 5 streams, indicating that evolutionary adaptation within taxa was also an important determinant of variation in this key trait among sites in our study. This finding is consistent with work demonstrating down-regulation of the metabolic normalisation in a unicellular alga via rapid (e.g. over 100 generations or 45 days) evolutionary adaptation to an experimental thermal gradient in the laboratory (Padfield *et al*. 2016). Collectively, this work highlights that changes in the metabolic normalisation result from temperature-driven selection both within and across species and can give rise to temperature invariance of metabolic rates along thermal gradients (Fig. 1b).

Our work shows that the mode of thermal adaptation, in driving complete temperature compensation of organism-level metabolism, had important implications for understanding the temperature dependence of ecosystem-level GPP across the catchment. GPP increased with temperature across the catchment (Fig. 3a), but it did so because biomass also positively covaried with temperature (Fig. 3b). After accounting for biomass, GPP was independent of temperature (Fig. 3c), consistent with the effects of thermal adaptation in driving temperature compensation of organism-level metabolism. These findings confirm the predictions of our model and previous suggestions (Kerkhoff *et al*. 2005; Enquist *et al*. 2007) that local adaptation of metabolic traits can yield the paradoxical phenomenon that rates of ecosystem metabolism are independent of temperature over thermal gradients that have been maintained over long timescales.

A great deal of empirical and theoretical work is still required to develop a complete, general theory that predicts how ecosystem properties emerge from evolutionary and community processes. Our work adds to recent efforts to this end (Enquist *et al*. 2007; Yvon-Durocher & Allen 2012; Schramski *et al*. 2015) by showing how temperature dependence of ecosystem biomass and the organism-level photosynthetic normalisation alter the emergent temperature sensitivity of ecosystem-level GPP. One important gap in the theory presented here is a mechanistic model for the temperature dependence of the metabolic normalisation owing to thermal adaptation. Our representation in equation 7 is merely a statistical description of an empirical phenomenon. The metabolic cold adaptation hypothesis seeks to explain the observation that species from cold environments often have higher mass-specific metabolic rates compared to counterparts from warmer regions as an evolutionary adaptation to compensate for lower biochemical reaction rates (Addo-Bediako *et al*. 2002). However, a quantitative, first principles derivation of this pattern remains elusive. Recent work on autotrophs has proposed that down-regulation of respiration rates as organisms adapt to warmer environments is driven by a necessity to maintain the carbon-use efficiency above a threshold when rates of respiration are more sensitive to temperature than those of photosynthesis (Padfield *et al*. 2016). Yet, as we have shown here, the assumption that the activation energy of respiration is always larger than that of photosynthesis does not always hold.

A better understanding of the mechanisms that give rise to the emergence of ecosystem properties is central to improving predictions of how global warming will alter the feedbacks between the biosphere on the carbon cycle (Levin 1998; Ziehn *et al*. 2011; Booth *et al*. 2012). Incorporating evolution into earth system and ecosystem models should be considered as a priority, especially in light of our finding that thermal adaptation can completely override the direct effects of temperature on metabolic rates. However, despite much recent progress (Smith & Dukes 2013; Daines *et al*. 2014; Smith *et al*. 2016), substantial work remains.

We capitalised on a ‘natural experiment’ using a geothermally heated stream catchment to show that thermal adaptation of photosynthesis drives an equivalence in biomass normalised GPP over a 20 ºC *in situ* temperature gradient. Our results suggest that local thermal adaptation plays a key role in determining how metabolic rates scale from populations to ecosystems and questions the assumption that the effects of temperature on enzyme kinetics can be applied to assess the long-term effects of temperature on ecosystem metabolism (Demars *et al*. 2016). They also shed light on the way in which the interplay between ecological and evolutionary processes could influence the response of the carbon cycle, and hence constituent food web and biogeochemical processes, to future environmental change.

## Acknowledgments

We thank Benoit Demars for providing reaeration data and comments that signficantly improved the manuscript. This study was supported a NERC Case studentship awarded to DP, GYD and SJ, an ERC starting grant awarded to GYD, and the University of Exeter.

## SUPPLEMENTARY INFORMATION

### Section 1. Supplementary Tables and Figures

**Figure S1.**
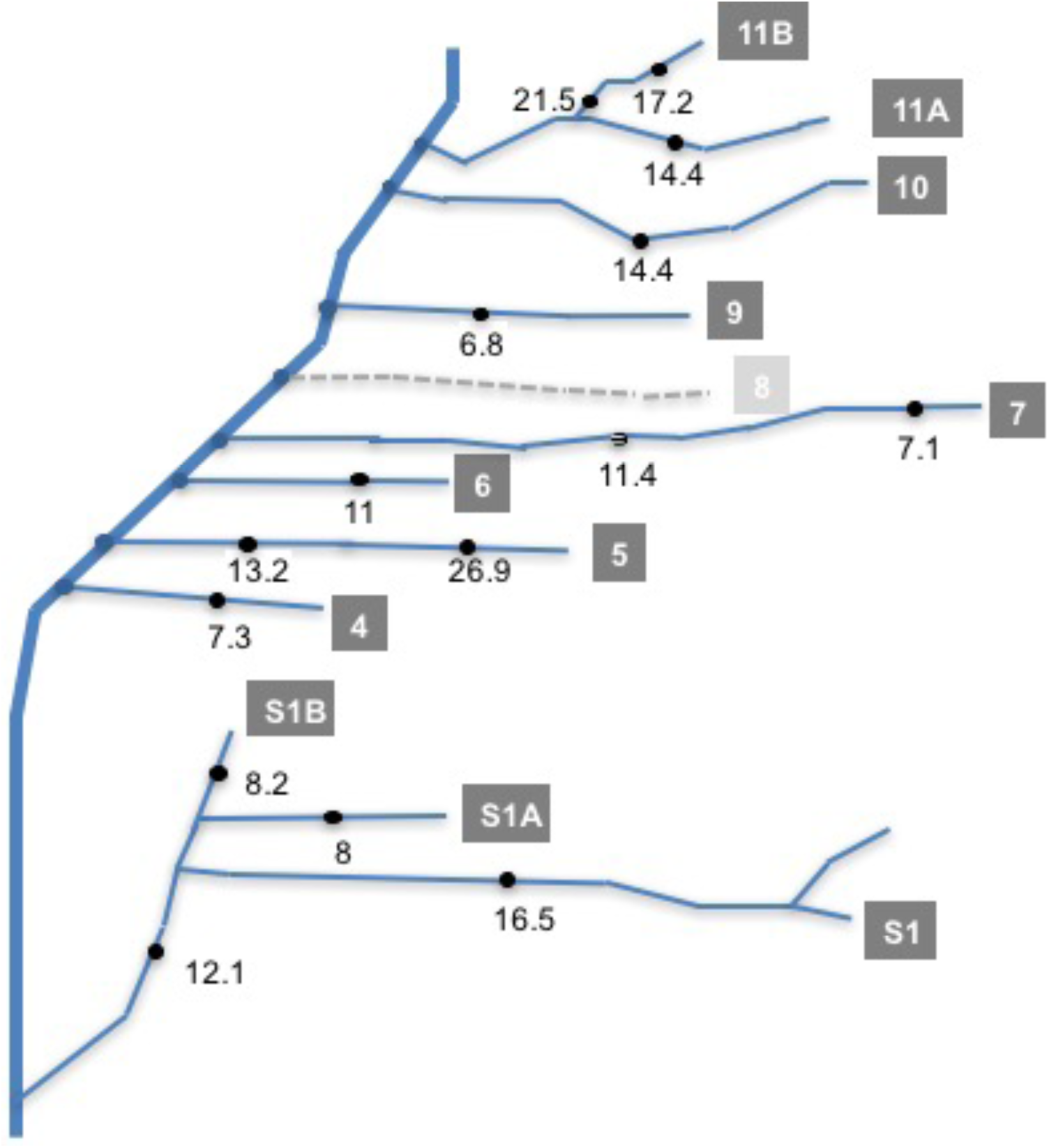
Map of the geothermal stream system in a valley near Hveragerdi, SW Iceland. Temperatures measured a various locations across the catchment are also given.

**Table S1.** Mean, minimum and maximum temperature values averaged across days and years (May 2015, May 2016) in the 15 sites. Values are based on a temperature estimates taken at 1 minute intervals. The streams are listed with increasing mean temperature.

| Stream | Temperature (°C) |  |  |
| --- | --- | --- | --- |
|  | Mean | Minimum | Maximum |
| S9 | 6.8 | 5.3 | 7.9 |
| S7 : high | 7.1 | 6.7 | 7.9 |
| S4 | 7.3 | 5.1 | 8.9 |
| S1A | 8 | 4.5 | 11.8 |
| S1B | 8.2 | 7.1 | 9.7 |
| S6 | 11 | 7.3 | 14.1 |
| S7 : low | 11.4 | 10.4 | 12.1 |
| S1 : low | 12.1 | 9.7 | 16.3 |
| S5 : low | 13.2 | 12.1 | 14.8 |
| S10 | 14.4 | 10.4 | 16.9 |
| S11A | 14.4 | 12.4 | 16.6 |
| S1 : high | 16.5 | 13.3 | 18.8 |
| S11B : high | 17.2 | 14.7 | 19.6 |
| S11B : low | 21.5 | 19.8 | 23.4 |
| S5 : high | 26.9 | 24.8 | 28.6 |

**Table S2.** Key physical and chemical features of the 15 sites investigated

| stream | width (m) | depth (m) | velocity (m s<sup>-1</sup>) | pH | conductivity (μS m<sup>-1</sup>) | nutrients (μmol L<sup>-1</sup>) |  |  |  |
| --- | --- | --- | --- | --- | --- | --- | --- | --- | --- |
|  |  |  |  |  |  | NO<sub>2</sub> | NO<sub>3</sub> | NH<sub>4</sub> | PO<sub>4</sub> |
| S9 | 0.41 | 0.027 | 0.11 | 7.57 | 173.3 | 0.29 | 0.23 | 0.27 | 0.86 |
| S7 : high | 0.4 | 0.053 | 0.3 | 7.43 | 359.1 | 0.22 | 0.44 | 0.28 | 0.7 |
| S4 | 0.46 | 0.06 | 0.36 | 7.27 | 204.6 | 0.2 | 0.08 | 0.22 | 0.14 |
| S1A | 0.59 | 0.07 | 0.5 | 7.40 | 230.9 | 0.25 | 0.4 | 0.7 | 0.54 |
| S1B | 0.42 | 0.058 | 0.14 | 7.50 | 462.4 | 0.28 | 0.25 | 0.18 | 0.17 |
| S6 | 0.19 | 0.029 | 0.12 | 7.43 | 289.6 | 0.22 | 0.4 | 0.21 | 1.02 |
| S7 : low | 0.3 | 0.043 | 0.4 | 7.43 | 304.7 | 0.22 | 0.44 | 0.28 | 0.7 |
| S1 : low | 1.1 | 0.13 | 0.81 | 7.36 | 305.2 | 0.26 | 0.26 | 0.48 | 0.35 |
| S5 : low | 0.32 | 0.041 | 0.09 | 7.63 | 273.6 | 0.22 | 0.57 | 0.17 | 0.14 |
| S10 | 0.22 | 0.109 | 0.24 | 7.53 | 167.0 | 0.35 | - | 0.24 | 0.74 |
| S11A | 0.71 | 0.078 | 0.77 | 7.17 | 235.7 | 0.24 | 0.29 | 0.19 | 0.55 |
| S1 : high | 0.74 | 0.12 | 0.61 | 7.20 | 321.7 | 0.26 | 0.26 | 0.48 | 0.35 |
| S11B : high | 0.31 | 0.042 | 0.33 | 7.33 | 407.9 | 0.25 | 0.25 | 0.27 | 1.25 |
| S11B : low | 0.4 | 0.042 | 0.33 | 7.33 | 407.9 | 0.25 | 0.25 | 0.27 | 1.25 |
| S5 : high | 0.17 | 0.037 | 0.06 | 7.63 | 319.2 | 0.22 | 0.57 | 0.17 | 0.27 |

**Table S3.** Pearson correlation coefficients between temperature and physical and chemical variables

| Variable | *r* | P value |
| --- | --- | --- |
| width | -0.14 | 0.56 |
| depth | 0.07 | 0.77 |
| velocity | 0.04 | 0.87 |
| pH | -0.03 | 0.91 |
| conductivity | -0.02 | 0.92 |
| NO<sub>2</sub> | -0.001 | 0.47 |
| NO<sub>3</sub> | 0.18 | 0.47 |
| NH<sub>4</sub> | -0.19 | 0.44 |
| PO<sub>4</sub> | 0.07 | 0.77 |

**Table S4.** The photosynthetic traits governing the thermal response curves for the 2 dominant biofilms of each stream.

| Stream | Year | Taxon | Net photosynthesis |  |  |  |  | Respiration |  |  |  |  | Gross photosynthesis |  |  |  |  |
| --- | --- | --- | --- | --- | --- | --- | --- | --- | --- | --- | --- | --- | --- | --- | --- | --- | --- |
| | | | $\ln np(T_c)$<br>($\mu\text{mol O}_2 \mu\text{g Chla}^{-1} \text{h}^{-1}$ @<br>10°C) | $E_{np}$<br>(eV) | $E_h$<br>(eV) | $T_h$ (°C) | $T_{opt}$<br>(°C) | $\ln r(T_c)$<br>($\mu\text{mol O}_2 \mu\text{g Chla}^{-1} \text{h}^{-1}$ @<br>10°C) | $E_r$<br>(eV) | $E_h$<br>(eV) | $T_h$ (°C) | $T_{opt}$<br>(°C) | $\ln gp(T_c)$<br>($\mu\text{mol O}_2 \mu\text{g Chla}^{-1} \text{h}^{-1}$ @<br>10°C) | $E_{gp}$<br>(eV) | $E_h$<br>(eV) | $T_h$ (°C) | $T_{opt}$<br>(°C) |
| S4 | 2016 | Cladophora | 3.9 | 1.03 | 4.39 | 30.6 | 28.48 | 2.48 | 1.01 | 3.78 | 31.66 | 29.53 | 4.2 | 0.92 | 9.19 | 32.49 | 30.57 |
| S1A | 2016 | Cladophora | 4.74 | 0.79 | 2.58 | 28.33 | 25.88 | 3.07 | 0.64 | 4.36 | 37.28 | 33.97 | 4.77 | 0.89 | 2.71 | 26.98 | 24.95 |
| S1A | 2016 | Nostoc | 4.37 | 0.52 | 8.77 | 36.87 | 34.28 | 3.14 | 0.44 | 4.26 | 38.83 | 34.63 | 4.8 | 0.47 | 2.72 | 35.36 | 30.71 |
| S4 | 2015 | Cladophora | 3.55 | 0.87 | 1.78 | 21.67 | 21.52 | 1.71 | 0.45 | 17.21 | 43.78 | 41.97 | 3.61 | 0.53 | 2.18 | 30.18 | 26.1 |
| S7 : high | 2016 | Cladophora | 4.35 | 0.73 | 7.04 | 31.77 | 29.33 | 3.85 | 0.8 | 2.94 | 31.06 | 28.42 | 4.48 | 0.93 | 3.51 | 28.24 | 25.99 |
| S7 : high | 2016 | Nostoc | 2.67 | 0.98 | 3.57 | 34.32 | 32.1 | 1.27 | 0.93 | 1.77 | 34.12 | 34.59 | 3.33 | 0.62 | 9.05 | 38.83 | 36.43 |
| S7 : high | 2015 | Feathermoss | 1.32 | 0.77 | 4.99 | 34.19 | 31.44 | 1.26 | 0.55 | 2.13 | 42.8 | 38.64 | 1.99 | 0.66 | 8.81 | 35.6 | 33.28 |
| S11A | 2016 | Nostoc | 2.67 | 1.91 | 5.29 | 28.47 | 27.62 | 1.13 | 0.71 | 5.85 | 45.74 | 42.79 | 2.95 | 1.57 | 4.37 | 28.46 | 27.43 |
| S10 | 2016 | Nostoc | 2.68 | 1.08 | 9.9 | 38.53 | 36.76 | -0.66 | 1.64 | 3.15 | 34.74 | 34.94 | 3.42 | 0.85 | 7.12 | 38.39 | 36.06 |
| S1 : high | 2016 | Nostoc | 3.77 | 1.03 | 8.87 | 39.61 | 37.69 | 1.33 | 1.09 | 3.23 | 41.01 | 39.26 | 4.36 | 0.85 | 3.66 | 37.86 | 35.15 |
| S11b : high | 2015 | Feathermoss | 1.82 | 1.12 | 2.64 | 24.53 | 23.64 | 1.1 | 0.48 | 1.64 | 49.7 | 45 | 2.08 | 1.14 | 2.5 | 25.19 | 24.64 |
| S5 : high | 2015 | Anabaena | 2.58 | 0.54 | 5.9 | 42.5 | 39.19 | 0.68 | 0.66 | 2.04 | 39.71 | 36.65 | 2.73 | 0.55 | 5.69 | 42.37 | 39.02 |
| S5 : high | 2016 | Anabaena | 2.66 | 0.85 | 4.02 | 37.4 | 34.71 | 0.45 | 1.58 | 2.35 | 29.63 | 32.12 | 2.79 | 0.77 | 5.63 | 39.89 | 37.14 |

**Table S5.** Results of a linear effects model analysis for each metabolic trait with fixed effects of stream temperature and metabolic flux (see Methods). Significant models are highlighted in bold.

| Metabolic Trait | Effect | d.f. | AIC | Log Lik | L-ratio | P value |
| --- | --- | --- | --- | --- | --- | --- |
| $b(T_c)$ | ~ 1 + stream temperature * metabolic flux | 8 | 96.94 | -40.47 | | |
|  | ~ **1 + stream temperature + metabolic flux** | 6 | 94.89 | -41.43 | 1.93 | 0.37 |
|  | ~ 1 + metabolic flux | 5 | 98.28 | -44.14 | 5.41 | **0.02** |
| $E_a$ | ~ 1 + stream temperature * metabolic flux | 8 | 37.03 | -10.51 | | |
|  | ~ 1 + stream temperature + metabolic flux | 6 | 36.36 | -12.18 | 3.33 | 0.189 |
|  | ~ 1 + stream temperature | 4 | 34.41 | -13.21 | 2.05 | 0.36 |
|  | ~ **1** | 3 | 32.92 | -13.46 | 0.51 | 0.48 |
| $E_h$ | ~ 1 + stream temperature * metabolic flux | 8 | 72.92 | -28.46 | | |
|  | ~ 1 + stream temperature + metabolic flux | 6 | 73.83 | -30.91 | 4.91 | 0.09 |
|  | ~ 1 + metabolic flux | 5 | 72.07 | -31.04 | 0.24 | 0.62 |
|  | ~ **1** | 3 | 71.37 | -32.68 | 3.30 | 0.19 |
| $T_h$ | ~ 1 + stream temperature * metabolic flux | 8 | -192.08 | 104.04 | | |
|  | ~ **1 + stream temperature + metabolic flux** | 6 | -192.92 | 102.46 | 3.15 | 0.206 |
|  | ~ 1 + metabolic flux | 5 | -190.32 | 100.16 | 4.60 | **0.032** |
| $T_{opt}$ | ~ 1 + stream temperature * metabolic flux | 8 | -27.21 | 21.61 | | |
|  | ~ **1 + stream temperature + metabolic flux** | 6 | -28.72 | 20.36 | 2.49 | 0.29 |
|  | ~ 1 + metabolic flux | 5 | -24.54 | 17.27 | 6.18 | **0.013** |
| $b(T_s)$ | ~ 1 + stream temperature * metabolic flux | 8 | 48.64 | -16.32 | | |
|  | ~ 1 + stream temperature + metabolic flux | 6 | 44.68 | -16.34 | 0.05 | 0.98 |
|  | ~ **1 + metabolic flux** | 5 | 42.99 | -16.49 | 0.31 | 0.58 |
|  | ~ **1** | 4 | 64.21 | -29.10 | 25.22 | **<0.0001** |

**Fig S2.**
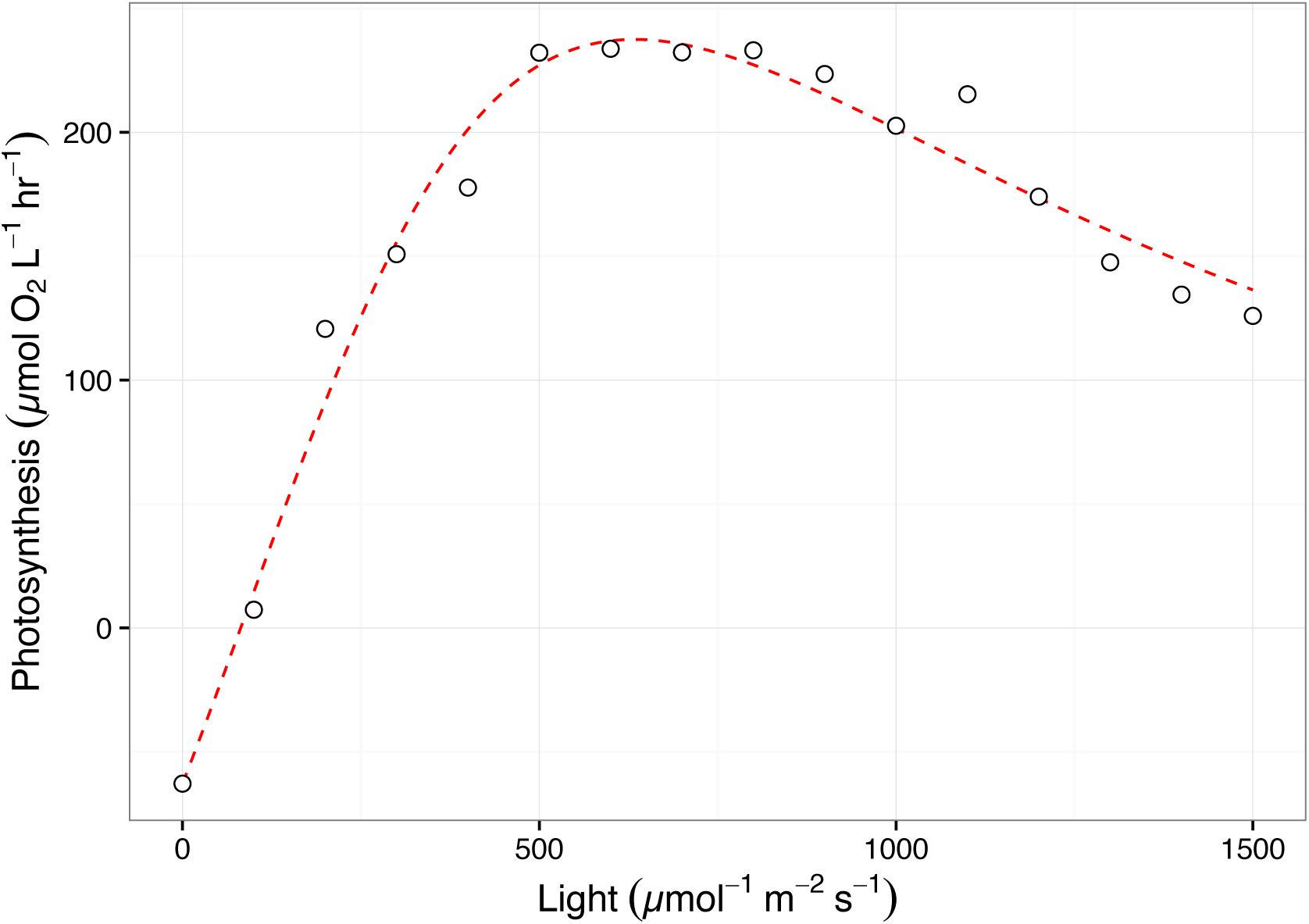
Photosynthesis irradiance curve used to determine optimal light for the acute temperature response of gross photosynthesis. Rates of net photosynthesis were measured at various light intensities at the average stream temperature of each biofilm. Here data are presented for *Nostoc spp.* in stream 7 (high) at 7.1 ºC. Lines represent the best fit to the modified Eiler’s model using non-linear least squares regression (See methods).

**Fig S3.**
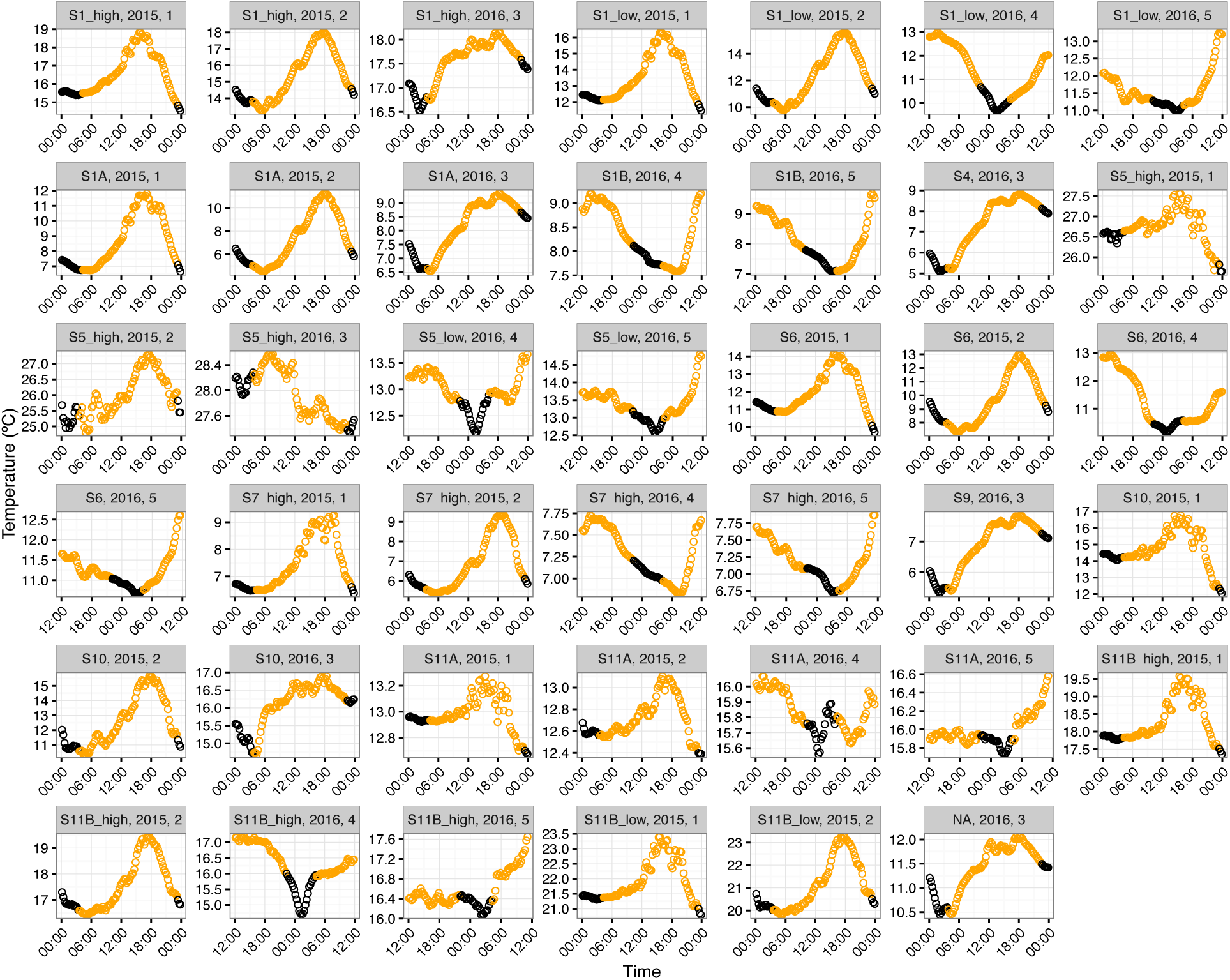
Daily cycles in temperature from each stream across days and years. Each panel is a single day of temperature variation split by each unique stream and across years (2015 or 2016). The data is split into “night” (black points) and “day” (yellow points) by defining night as < 5µmol m^-2^ s^-1^ (see Methods).

**Fig S4.**
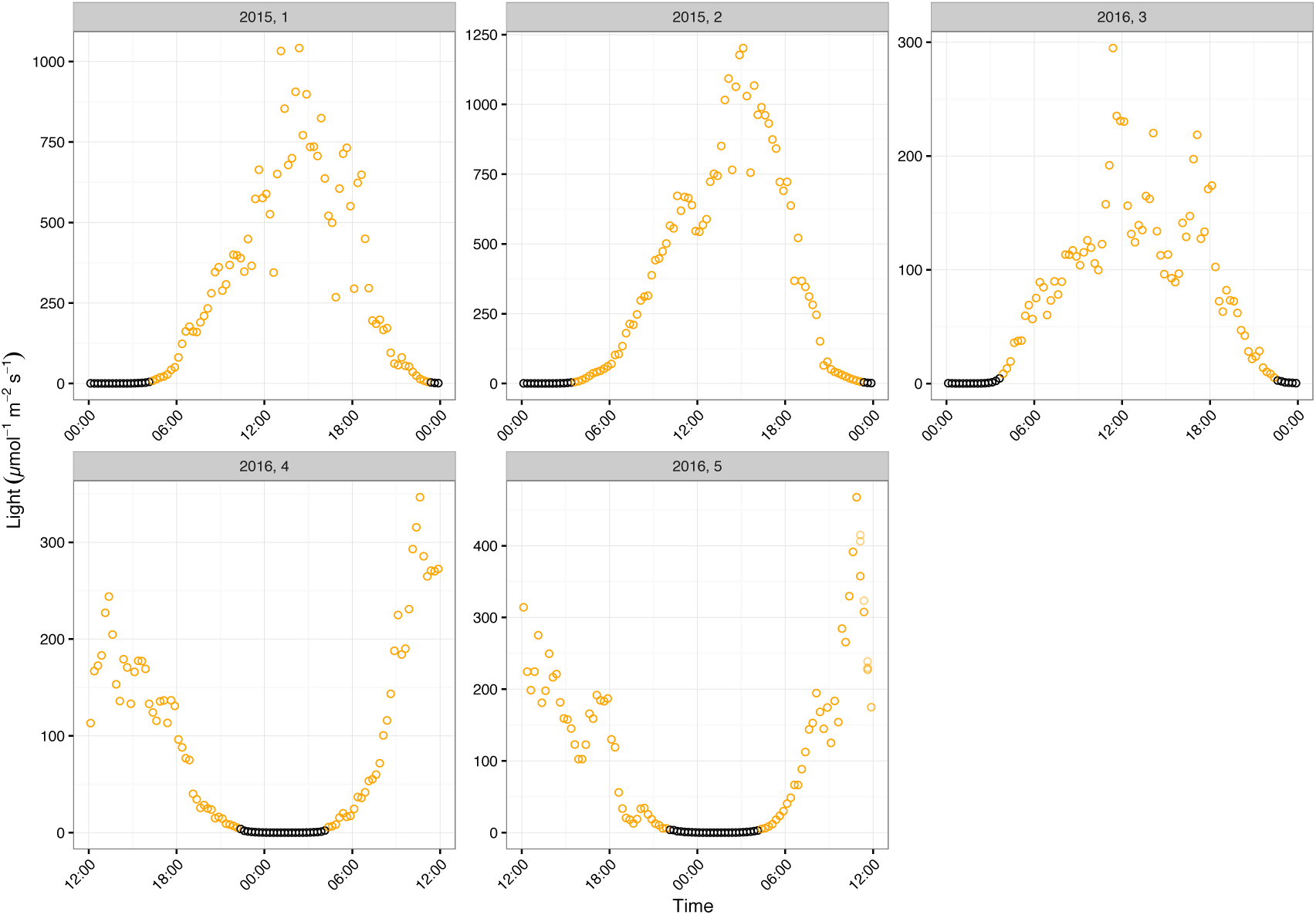
Daily cycles in light from across days and years. Each panel is a single day of light variation split by each unique stream and across years (2015 or 2016). The data is split into “night” (black points) and “day” (yellow points) by defining night as < 5µmol m^-2^ s^-1^ (see Methods).

**Fig S5.**
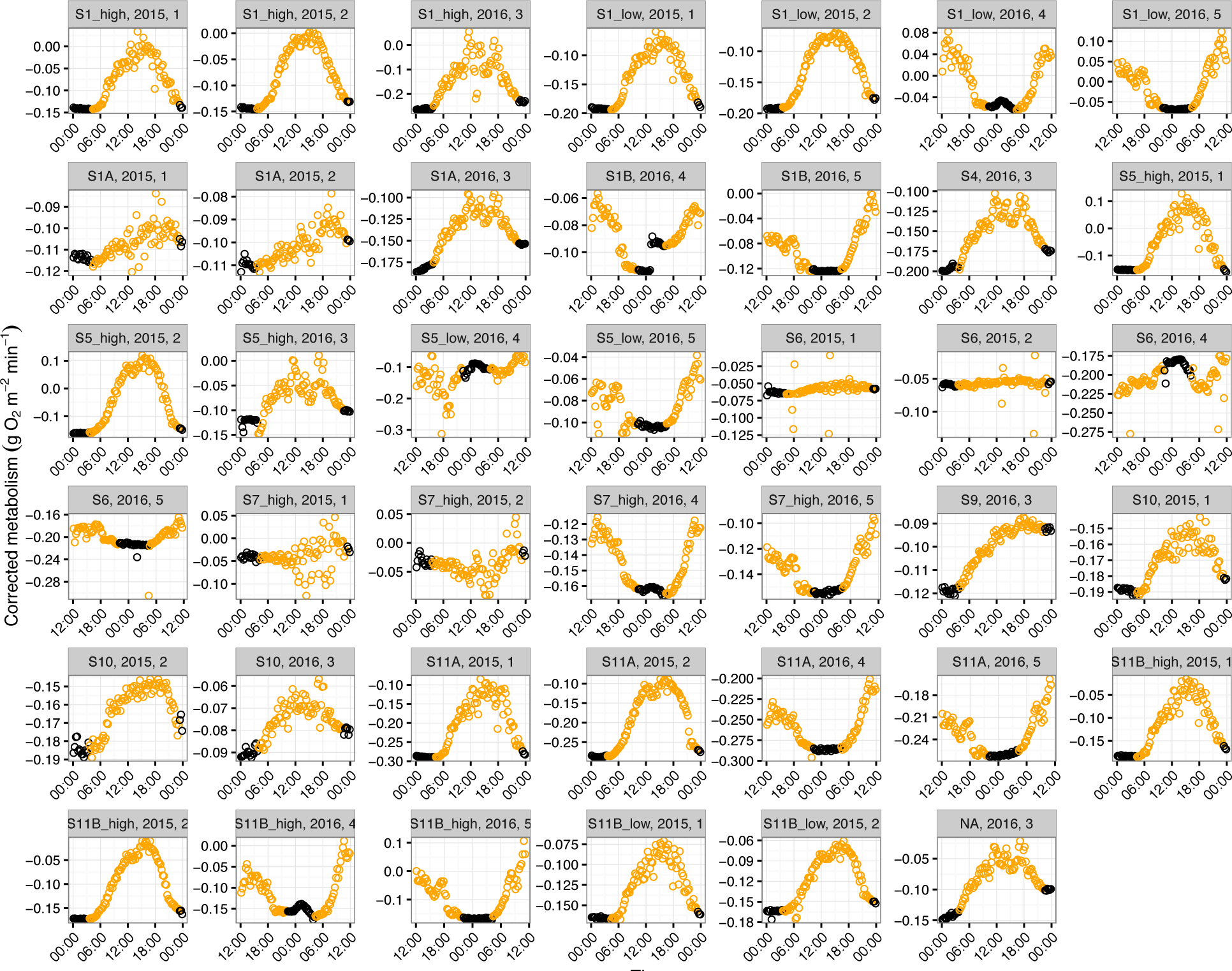
Daily cycles in metabolic flux from each site across days and years. Each panel is a single day of metabolic rate after accounting for reaeration (Δ*DO – K*; see Methods) split by each unique stream and across years (2015 or 2016). The data is split into “night” (black points) and “day” (yellow points) by defining night as < 5µmol m^-2^ s^-1^ (see Methods).

**Fig S6.**
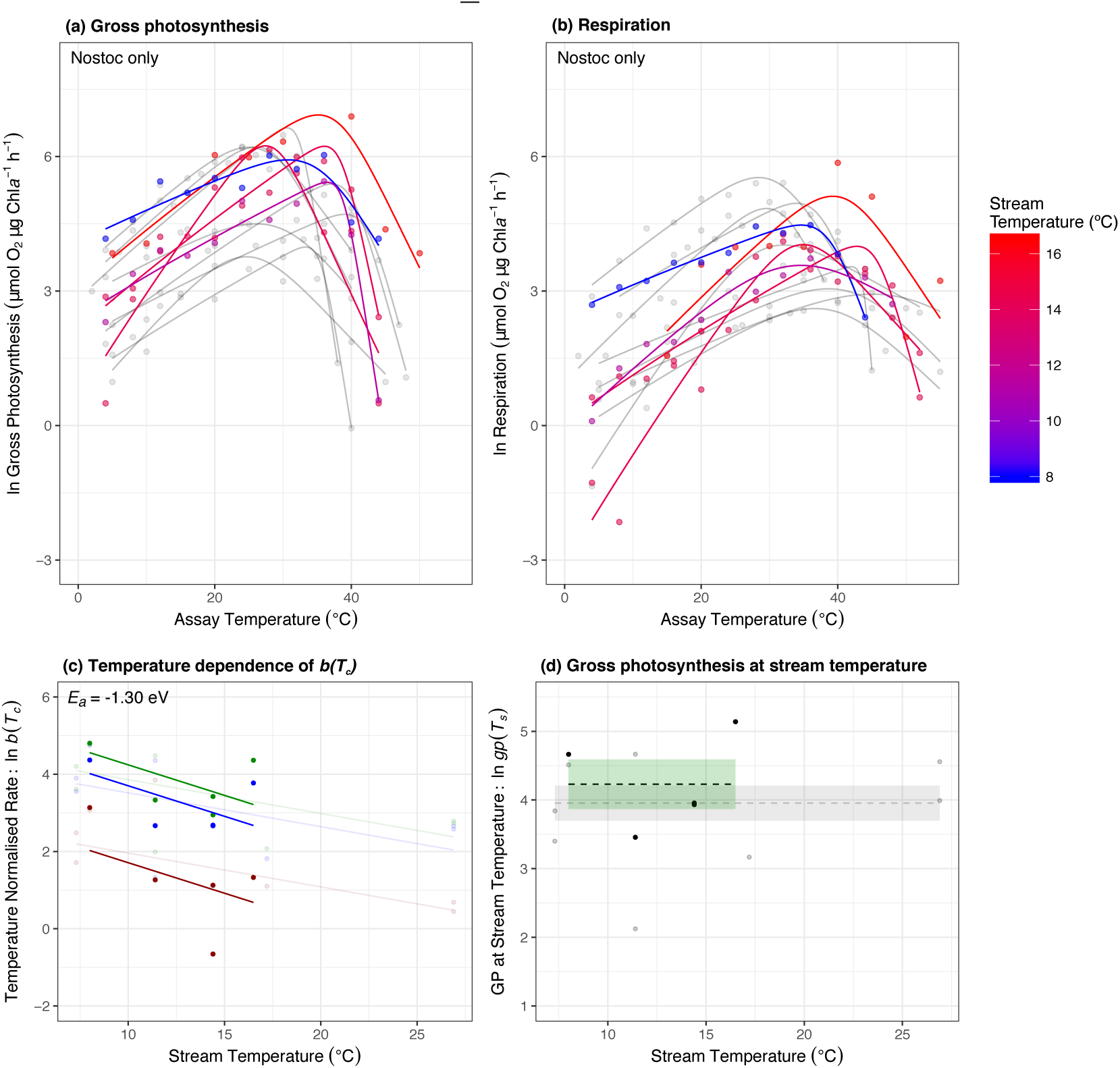
Patterns of thermal adaptation in *Nostoc* spp. only. (a) (a, b) Acute thermal response curves for gross photosynthesis and respiration were measured for each isolated autotroph from streams spanning average temperatures from 7 ºC (blue) to 17 ºC (red) for stream biofilms dominated by *Nostoc* spp. (c) Optimum temperatures were consistently higher than the average stream temperature. (c) Metabolic rates normalised to 10 ºC,b(T_c_), decrease exponentially with increasing stream temperature for gross photosynthesis (green), net photosynthesis (blue) and respiration (red). (d) Rates of gross photosynthesis at the average stream temperature showed no temperature dependence. Grey points and lines highlight the other taxa to facilitate direct comparison to the relationship for *Nostoc* spp.

### Section 2. Supplementary Methods

**Derivation of the activation energy of net photosynthesis.** The rate of net photosynthesis, *np*(*T*), at temperature, *T*, is equal to the difference between the rates of gross photosynthesis, *gp*(*T*), and respiration, *r*(*T*). Equation 5 (main text) implies that the temperature sensitivity of net photosynthesis will not follow a simple Boltzmann-Arrhenius relationship. Instead, the apparent activation energy of net photosynthesis, *E_np_*, can be approximated in the vicinity of *T_c_* as (Yvon-Durocher *et al*. 2014),

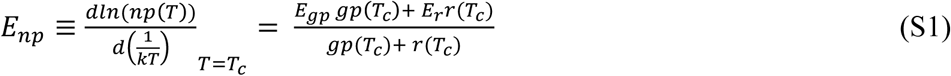
which is equal to an average of the activation energies of *E_gp_* and *E_r_*, weighted by their respective normalisations, *gp*(*T*) and *r*(*T_c_*) Using this approximation, we can then express the temperature dependence of *np* as

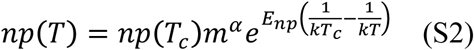
where *np*(*T_c_*)= *gp*(*T_c_*)− *r*(*T_c_*). We quantified the accuracy of this approximation by comparing *E_np_*, derived using eq. S1 to the apparent activation energy of net photosynthesis measured by fitting eq. (1) to the net photosynthesis data (see Methods). The derived and measured estimates of *E_np_* were positively correlated with a slope that had confidence intervals which overlapped unity (slope = 1.22, 95% CI: 0.78 – 1.65) and *R*
^*2*^ = 0.75 (Fig. S7).

**Figure S7.**
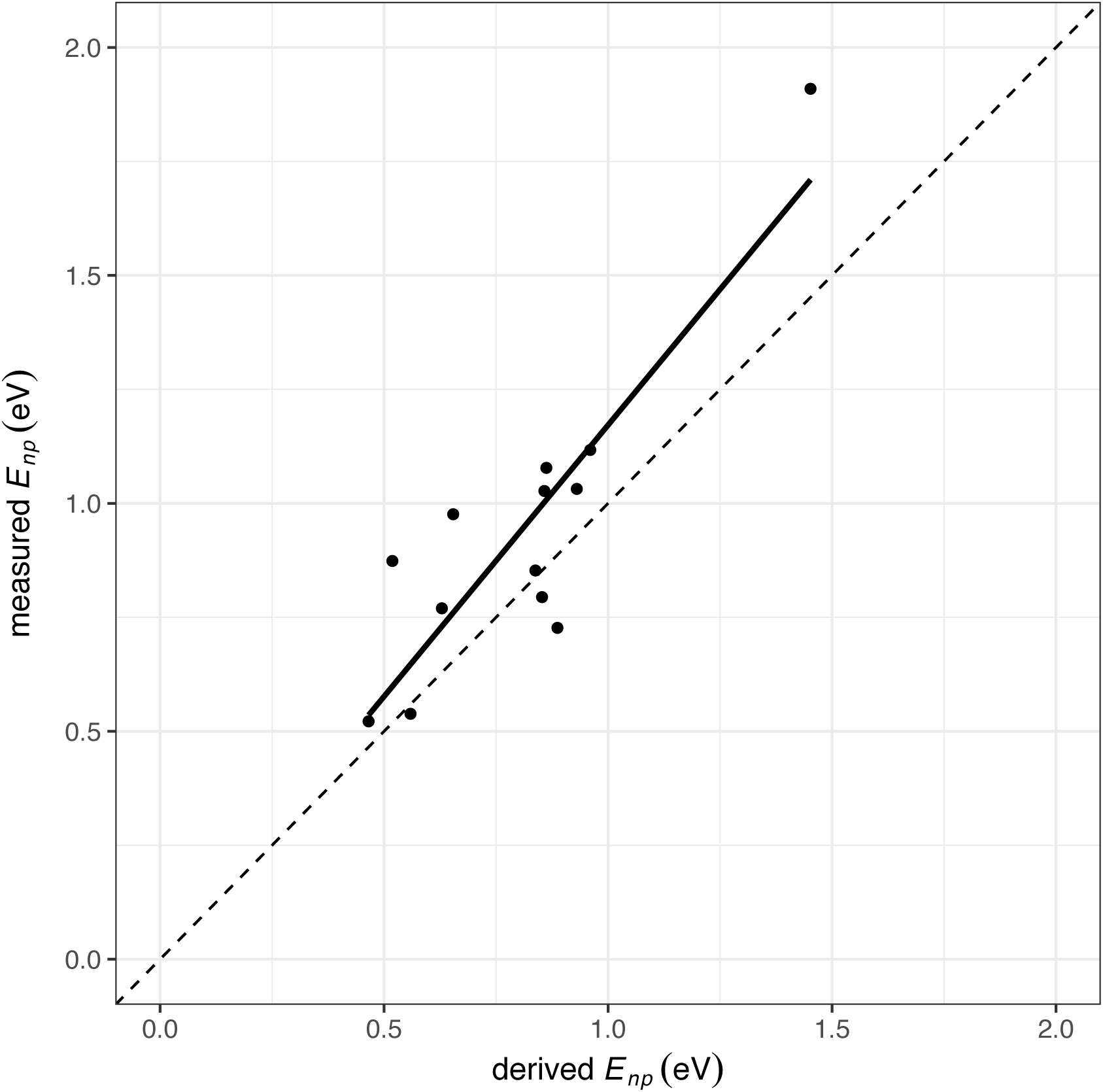
Comparison between measured and derived activation energies for net photosynthesis. Activation energies of net photosynthesis measured from fitting the rate data to the modified Sharpe-Schoolfield equation (eq. 1) correlate well with the derived activation energy of net photosynthesis calculated using equation S1. The fitted line is the best fit of a linear model and the 1:1 line is shown for comparison.

**Comparison of measured and modelled reaeration rates.** To assess the robustness of our modelled values of reaeration, we compared measurements of the reaeration rate made in nearby streams in Iceland with comparable physical characteristics using propane additions (from Demars *et al*. 2011), to values estimated using the surface renewal model (eq. 14, main text). In Demars *et al*. (2011), the reaeration rate was measured using a tracer study, where propane was bubbled continuously across the width of the stream at an upstream station. Water samples were taken at a downstream station and analysed by gas chromatography back in the laboratory (see for a more detailed description of the methods). The change in propane concentration the over the reach and the travel time were used to estimate the reaaeration rate, *K* (min^-1^).

We compared the measured values of reaeration, *K* (min^-1^), from Demars *et al*. (2011) to estimated values of *K* derived Eq. 14 (main text) and measurements of velocity, depth and temperature for those streams. We found a strong correlation between modelled and measured values of *K* with 95% confidence intervals on the slope that included unity (slope = 1.13, 95% CI: 0.76 – 1.50) and an *R^2^* = 0.61 (Fig. S8). Consequently, we are confident that estimates of reaeration derived from the surface renewal model are robust for the streams included in our survey.

**Figure S8.**
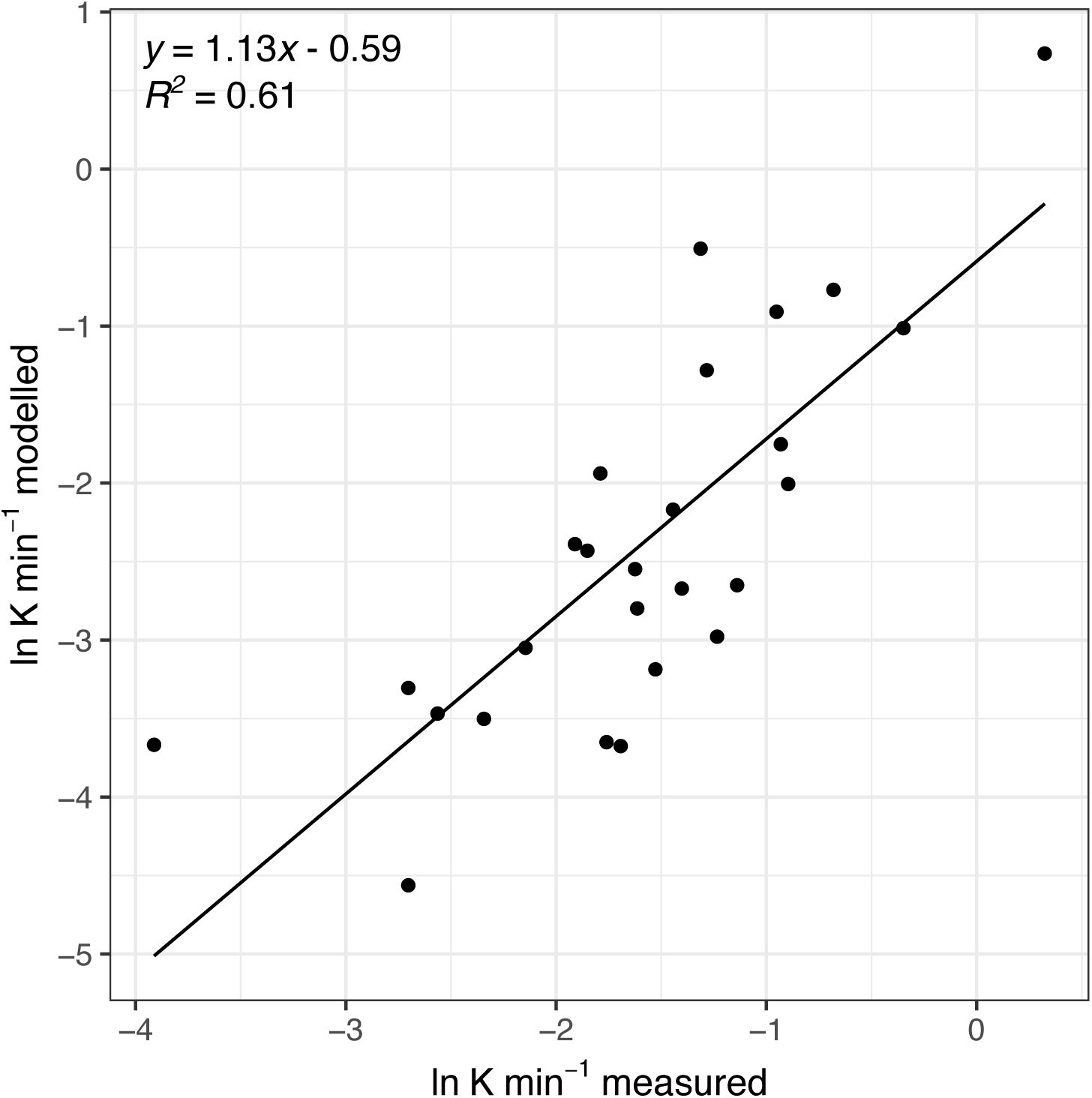
Comparison of modelled and measured rates of reaeration. Rates of measured reaeration using a propane tracer study are positively correlated with those derived using the surface renewal model (eq. 14; main text) with slope that was statistically indistinguishable from unity.

